# Trust in scientists and rates of noncompliance with a fisheries rule in the Brazilian Pantanal

**DOI:** 10.1101/468728

**Authors:** Ethan A. Shirley, Meredith L. Gore

## Abstract

Natural resource rules exist to manage resources and the people that interact with them. These rules often fail because people do not comply with them. Decisions to comply with natural resource rules often are based on attitudes about legitimacy of rules and the perceived risks of breaking rules. Trust in agencies promulgating rules in part may determine perceptions of legitimacy of the rule, and in turn depends on individuals’ trust in different agency actors. The purpose of this research was to explore the relationship between fishing rule noncompliance and trust in scientists, a key group within management agencies. We interviewed 41 individuals in one rural fishing community in the Brazilian Pantanal from April to August, 2016, to assess (1) noncompliance rates, (2) noncompliance-related attitudes, and (3) the relationship between trust in scientists and noncompliance decisions in the region. We found that among study participants, noncompliance was common and overt. Trust in scientists performing research in the region was the best predictor of noncompliance rate with a fishing rule (nonparametric rank correlation *ρ* = −0.717; Probit model pseudo-R^2^ = 0.241). Baseline data from this research may help inform future interventions to minimize IUU fishing and protect the Pantanal fishery. Although our results are specific to one community in the Pantanal, trust in scientists is potentially an important factor for compliance decisions in similar situations around the world. These results build not only on compliance theory but also speak to the important role that many scientists play in the geographic areas in which they conduct their research.

## Introduction

As human populations grow, they can increase pressure on the environment in which they live and the natural resources on which they rely (1,2). Environmental rules—such as laws, regulations, and social norms—exist to help mitigate risks associated with anthropogenic pressures. Unfortunately, the rules that exist to ensure the persistence of natural resources often fail to do so fully. Natural resource rules usually fail in one of two ways: they are poorly defined (i.e., even if everyone follows the rule, the natural resource will be exhausted because limits are inadequate); or, they are well defined but not followed (i.e., people do not always comply with the rule). Research on compliance and noncompliance therefore is important to examine failures of rules to manage human pressure on the environment. Oftentimes researchers and practitioners work to address noncompliance and compliance concomitantly (3–5). These dual-mission efforts continue despite recognition that within the context of conservation, motivations for compliance are not necessarily the inverse of those for noncompliance, or the violation of rules (6). Irrespective of divergent motivations of noncompliance and compliance, however, decreasing rates of intentional noncompliance can help overcome the second type of rule failure.

The case of illegal, unregulated, and unreported (IUU) describes failures of natural resource rules either by fishing in violation of them or in their absence. IUU fishing poses risks to fisheries and humans worldwide (7). In particular, IUU fishing poses risks to the size and composition of fish populations from overharvesting, risks to ecosystem health and function from degradation, and risks to humans from reduced income from tourism and professional fishing (7). This is significant because fisheries play a foundational role in sustaining healthy ecosystems and providing food security for billions of people worldwide (8). IUU fishing is increasingly recognized as a global high policy priority issue, with the United Nations, civil society groups, nongovernmental organizations, and governments working, often together, to reduce its associated risks to both global fisheries and the billions of people that depend on them (9). Reducing noncompliance and thereby increasing rates of compliance, which is unintentional or intentional behavior in adherence with laws and rules (5), is one mechanism for reducing risks from IUU fishing.

The extant literature includes foundational insight into many answers for questions underlying noncompliance with IUU fishing. In a marine context, higher levels of risk of getting caught by surveillance can increase compliance rates by decreasing noncompliance with rules (10). However, surveillance and policing in rural and remote areas is often difficult and costly. Other lines of inquiry demonstrate perceived legitimacy of rules and rule makers are important factors influencing decisions to intentionally comply or not comply with laws (11–14). Attitudes about legitimacy can be intertwined with perceived risk (15). Risks to the environment can difficult for individuals to assess, and often perceptions of risks and causes of environmental degradation differ considerably between laypeople, rule makers, and scientists involved in setting rules. Questions remain about the suite of attitudes underlying individuals’ decisions to comply or not comply with conservation-based rules. This gap in understanding widens when questions about compliance and IUU fishing are considered within the inland, freshwater fishing context. Inland fishing contexts may present distinct challenges from marine fisheries because they represent restricted habitats that are easier to access by private parties than many areas of the open ocean. The few studies that do focus on this area avoid inquiry about perceived environmental risks and legitimacy of rules (16). One meaningful gap between risk perception and legitimacy is trust, including trust in individuals associated with rulemaking agencies and the institutions that these individuals represent. Agencies and politicians are often geographically far-removed from the natural resources they are responsible for managing, while scientists often work directly with natural resources and in the communities that use those resources (17). Trust in scientists may therefore represent an important part of certain people’s noncompliance or compliance decisions.

In this work, we consider the case of inland IUU fishing in one community in the Brazilian Pantanal. In the Pantanal, a key region for conservation of biodiversity, scientists’ research in rivers helps inform legal limits for fishing. At the same time, trust in science is thought to be decreasing (18,19). Our first objective was to assess noncompliance rates in the region. Our second objective was to gauge attitudes about risk and natural resource management in the region. Our third objective was to explore the relationship between trust in scientists, risk perception, and noncompliance. Our interdisciplinary approach reflects that of conservation criminology, or the integration of natural resource management, criminology, and risk and decision science (20). Enhanced knowledge about why people choose to violate rules can inform the design and evaluation of crime prevention programs and policies as well as law enforcement monitoring (6). The primary aim of this work is to build new knowledge that advances interventions to reduce IUU fishing in the Pantanal and help minimize risks to the fishery and people that interact with it.

## Conservation criminology: risk, trust, and natural resource management

Conservation criminology, as the science of conservation crime, uses insights from the fields of risk and decision science, natural resource management, and criminology (20).This interdisciplinary approach offers one lens to understand human behavior associated with illegal natural resource use. Conservation criminology advises consideration of criminology to understand conservation behavior and violations of conservation rules. Criminologists characterize intentional compliance with rules as being either coerced or voluntary. Coerced compliance generally relies heavily on policing and penalties for offenders (21,22), and it is on the manner of coercion (e.g., increasing detection or punishment; 23) that many criminologists focus. These coercion-based compliance studies look at external controls of behavior through fines and jail time for offenders who are caught (10, 21–23). Theoretically, people who calculate the risk of getting caught as being too high and the punishment too severe are deterred from engaging in noncompliant behavior (21). However, IUU fishing often occurs in regions where rule enforcement is not economically or physically viable. For example, areas in the middle of the open ocean can simply too vast to patrol closely and inland lakes can be surrounded by forests with unreliable ports of entry, inaccessible roads or other ingresses. In some instances, private landowners shield offenders from law enforcement authorities. Where coerced compliance is not viable, the natural resource management and risk and decision science parts of conservation criminology are especially valuable analytical tools.

Voluntary compliance is not coerced; instead, this type of compliance results from individual decisions to follow, rather than break, the rules, and has been the focus of more recent compliance work in the natural resources context (13). Approaching noncompliance with IUU rules through the lens of risk and decision science offers one way of studying voluntary compliance among individuals. Decisions to comply with or violate rules can be thought of as individuals’ cost-benefit analyses, with costs differing depending on views about agency actors, the rules, and the environment itself cites. Behavioral decisions can be influenced by attitudes (see 24), and many attitudes are themselves influenced by the structure of natural resource management. Attitudes about fisheries conservation rules, including trust and legitimacy, can influence individuals’ responses to those rules (25). Attitudes affect perceptions of risk (i.e., external cues are utilized based on internal attitudes) (26). Decisions under uncertainty are fundamentally different than cognitively simpler decisions with clear costs and benefits (27,28). In this instance, risk can be defined as the probability and the negative value—damage, associated with an action (29). Risk perception generally describes the intuitive judgments people make about risks as opposed to the technical assessments made by experts (30). Environmental risks can be particularly difficult to assess in decision-making processes because they are often uncertain and difficult to quantify (31). When people individually make decisions to harvest common pool natural resources such as fish, the damage they theoretically perceive themselves causing to the resource (i.e., the risk) is a fraction of the gain that they personally receive (1). Rules help clarify the acceptable levels of environmental risk, thus facilitating decision-making by identifying and quantifying damage that might otherwise not be readily apparent (31).

Finally, conservation criminology requires considering the natural resource dimensions of IUU fishing. Natural resource management (NRM) authorities, such as government agencies, help clarify risks by promulgating environmental rules. Empirical studies place attitudes relating to legitimacy of rules and of rule makers among the range of attitudes influencing compliance with laws (11,15,32).We know legitimacy is related to trust, or the willingness to accept vulnerability (33), and perceived procedural fairness in a NRM authority. Trust, perceived procedural fairness, and legitimacy have been suggested to affect compliance decisions (34,35). Trust in agencies is in part a function of trust in agents of the agency or rule makers as individuals (33). Trust depends in part on trustworthiness factors grouped by some authors into categories of identity, ability, benevolence, and integrity (36,37). Others have analyzed perceived procedural fairness, or fairness of the procedures behind creation and enforcement of laws and rules, separately (38). The questions used in the literature to measure trust, procedural fairness, trustworthiness, and legitimacy are similar (39). Maximizing positive NRM outcomes such as successful sustainable use can be associated with increased or maintaining trust in management authorities (40). Trust helps explain why community-based natural resource management (CBNRM) can lead to more enduring, sustainable, and publicly accepted conservation outcomes over top-down natural resource decision-making by federal or state agencies (41). Conversely, lack of trust in natural resource authorities and agency contribute to delegitimizing the protective conservation measures promulgated by agencies, including rules. Without legitimate rules from trusted NRM agencies, people may perceive environmental risks differently than the agencies and be less consistent in their voluntary compliance.

The compelling relationship between trust in agencies and positive natural resource management outcomes has been explored in many different conservation contexts (33,34,42). Interestingly, although natural resource management occurs at different geographic scales (e.g., local, national, transfrontier), trust is often measured at a single scale: managers and management (35). It is noteworthy then that studies exploring the relationship at the local scale, or between trust in scientists and noncompliance, do not exist in the literature, because many scientists do their work in the field often in or near communities impacted by natural resource rules. Considering the influence of trust at different scales may be especially important where rule makers are seen as outsiders imposing rules from a distant capital; considering trust in scientists, specifically, as part of the rulemaking authority, may be especially important in areas where scientists are actively and visibly involved in research. This situation is common in certain rural communities where scientists doing research on natural resources are seen as the local arm of power-wielding agencies (17). To this end, we framed our exploration of trust and compliance at the level of the scientist.

## Study area: the Brazilian Pantanal and its fisheries management context

The Pantanal is among the world’s largest wetlands (43), spanning 150,000 square kilometers in the center of South America and stretching over parts of Bolivia, Paraguay, and Brazil (Fig 1). The largest proportion of the Pantanal belongs to Brazil, where its rivers, lakes, forests, and savannas provide refuge for endangered species of fauna and an important migration stop for birds. The Pantanal drains part of the central Cerrado high plains of Brazil and its rivers feed into the De La Plata River basin before emptying into the Atlantic Ocean near Buenos Aires and Montevideo. The Pantanal is recognized as a key biodiversity area because of the role it plays in regional hydrology, collecting, filtering, and funneling water into the Paraguay-Paraná River system (44). It is also recognized as a key conservation area for its rich biodiversity, including endangered and threatened species like the hyacinth macaw (45). Despite its priority status, in Brazil the Pantanal’s lands are over ninety-percent privately owned (46); thus, private citizens’ compliance with existing environmental laws and rules is critical to its conservation. Thousands of people live in the Pantanal, sparsely distributed over the vast, seasonally-flooded mosaic of forests, rivers, and savannas. Enforcement efforts to maximize compliance with comprehensive environmental regulations are hampered by a lack of infrastructure, and their efficacy is not well understood because patterns of and motives for noncompliance have never been studied in the region.

**Fig 1.**
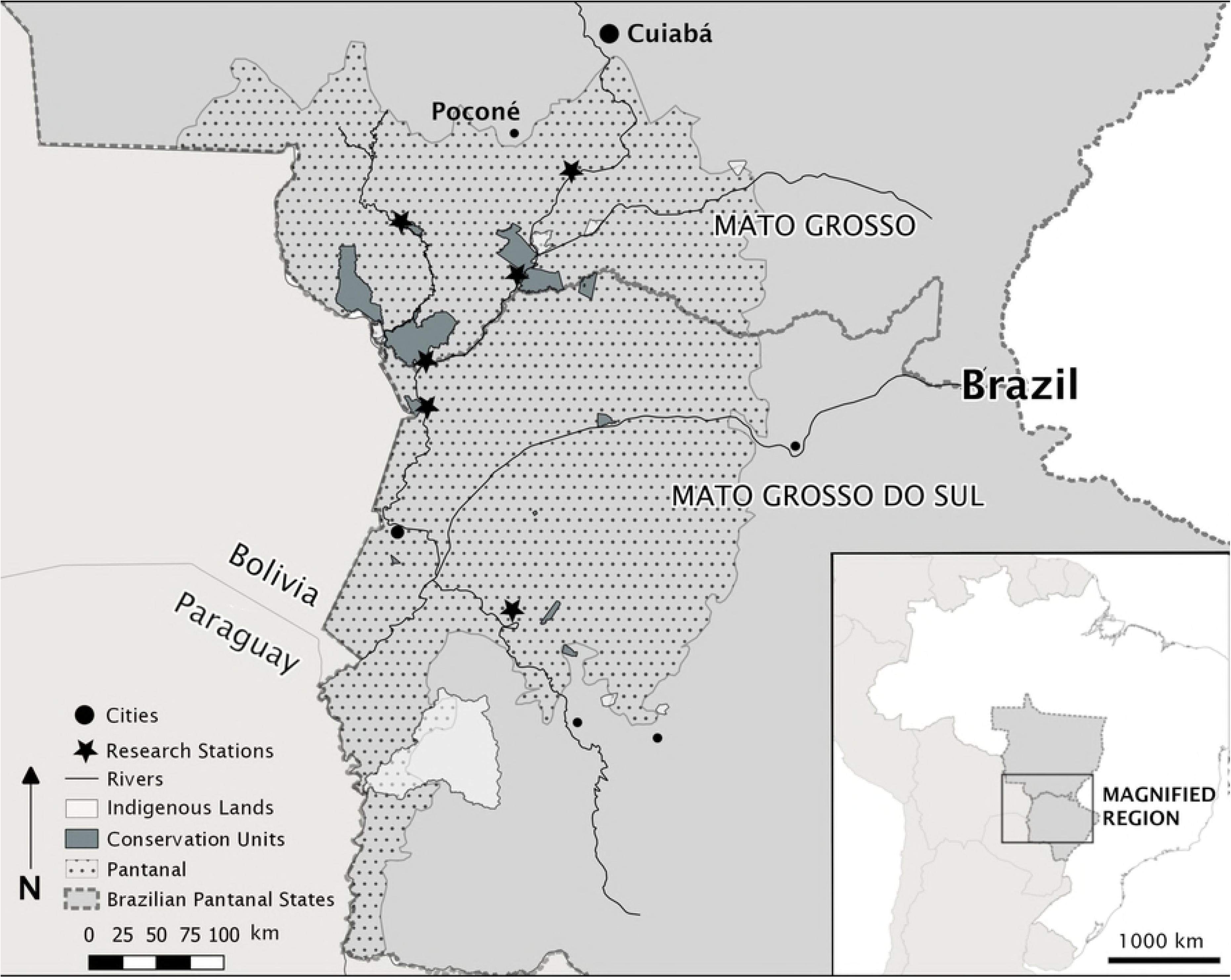
Map of the Brazilian Pantanal and research stations in regional context. The Brazilian Pantanal occupies parts of Mato Grosso and Mato Grosso do Sul within Brazil, and borders Pantanal regions in Bolivia and Paraguay. Cities, towns, conservation units, lands set aside for use by indigenous peoples, and approximate locations of some research stations within the Pantanal. The community in this study is located outside of Poconé, some 150km from Cuiabá, where monthly rulemaking meetings take place.

Conservation challenges in the Pantanal include IUU fishing (47). There are three types of regulated fishing in the Pantanal: amateur, subsistence, and professional-artisanal (also called just “professional”). Fishermen are organized into municipal fishermen’s colonies, which function as an advocacy-type lobby representing fishermen’s rights in each municipality (48). Many people who work as professional fishermen live in areas that are largely inaccessible to the relatively small number of enforcement officers who have limited patrol resources and basic levels of policing technology. In this regard, individual voluntary compliance with rules especially important in the Pantanal (47). The organ responsible for setting the fishing rules for all types of fishing in each state is called the Fishing Council, (*Conselho da Pesca*, or CEPESCA), which involves a mixture of top-down and participatory co-management. In Mato Grosso, it is composed of scientists from the local state and federal universities, representatives from regulators at the State Secretary of the Environment (SEMA), and members of fishermen’s colonies, along with legislators. CEPESCA defines laws and rules based on scientific research and the needs of fishermen and other community members, who are free to contribute to public debates and focus groups with legislators and others who draft the rules. The primary market fish in the region are three siluriforms (catfish) and four characiforms (piranha-like fish), including the pacu (*Piaractus mesopotaminus*) (47). CEPESCA regulates fishing in the region by creating a minimum size limit for each species and a weight limit depending on what type of fishing license fishermen possess (49).

## Methods: participants, instrument, and analysis

### Case study respondents

We focused our inquiry on *in-loco* professional fishermen in the municipality of Poconé. *In-loco* professional fishermen in the region are a key stakeholder group with a vested interest in preserving the environment of the Pantanal for sustainable use. These professional fishermen live permanently on the banks of rivers and have for generations, and therefore have longstanding ties to the land and the sustainable harvest of resources in the region (50). Previous work with local fishermen sought to representatively sample the fishermen’s colony as a single stakeholder group (48). However, as many as two-thirds of all professional fishermen live in cities and use their professional license to collect welfare during the spawning season when fishing is closed (51). We distinguished these two groups because of the possibility of their having different incentives to conserve the fishery—*in-loco* stakeholders have diverse ties to local natural resources that extend beyond the purely monetary.

The group of respondents for this study consisted of all the active professional fishermen belonging to Colony Z-11 living in one port community along the Cuiabá River in the Poconé municipality in between April and August, 2016. The community is sparsely distributed and not well delimited, so we considered for this study only the most densely populated region one-hour by speedboat upriver and downriver of the port. The lead author, fluent in Portuguese, visited every domicile and interviewed everyone found living on that part of the river and over the age of 18; in this regard study respondents represent a good faith and complete subset of *in-loco* professional fishermen living in the community during the study period.

### Instrument design and implementation

Our first objective was to assess noncompliance rates. Interview questions asked directly about people’s perceptions of others’ noncompliance rates in the community as well as their own noncompliance rates with a specific rule that was universally known to fishermen in the region. Second, we focused on exploring the factors underlying noncompliance. We asked direct questions about why people think other people choose to violate rules. Then, we assessed attitudes about risk and trust as factors that impact noncompliance decisions. Attitudinal and risk questions were taken from the English literature, translated into Portuguese by the lead author and pretested with fishermen (n = 7) for construct validity and ease of understanding before they were included in the survey instrument. Trust and trustworthiness questions were replicated from (34) and (33), as well as (37). Questions were selected to represent aspects of trust and trustworthiness that elsewhere in the literature have been called procedural fairness (32,34). Environmental risk questions were derived from (52).

We used a voluntary questionnaire verbally administered face-to-face because most individuals within the target population were not literate and did not have reliable access to mail, internet, or land-line phones. The survey instrument began with a statement informing participants of the intent of the research, including ensuring participant confidentiality and researcher independence to mitigate effects of bias in responses (53). Following the statement of informed consent, we asked first general questions focusing on environmental attitudes following Gore et al. (52). We followed these questions with projective questions about noncompliance rates and reasons (e.g., asking individuals to describe incidences of other people’s noncompliance). Then, we asked a prospective question about noncompliance (i.e., inquiring about possible individuals’ future rates of noncompliance). Both projective and prospective questions about noncompliance have been shown to reduce bias in responses about noncompliance (54). The single question about prospective personal noncompliance was placed at the end of the interview to minimize the effects of the social desirability bias (55).

Demographics were assessed following the completion of the substantive parts of the survey. The survey took approximately ten minutes to administer. All subjects’ identities were protected and we did not ask their names. Michigan State University’s Institutional Review Board, specifically the Human Subject Protection Program, was the ethics committee that reviewed and approved these methods exempt from review for the duration of the research (IRB x15-643e).

### Measurement and data analysis

Attitudinal questions were measured on a five-point Likert-type scale (1 = “Disagree Completely” to 5 = “Completely Agree”). Noncompliance was assessed with a five-point frequency question (1 = “Never,” 2 = “Rarely,” 3 = “Sometimes,” 4 = “Often,” 5 = “All the time”), following questions asked in (56). For our first objective, we report proportions of each response to questions about community noncompliance rates and some notable correlations. For our second objective, we report proportions of each response to questions about perceptions of motivations behind noncompliance as well as means, medians, and standard deviations for attitudes and risk perceptions. A composite score of responses to trustworthiness and procedural fairness questions was created using the mean of responses. We report relevant Spearman’s *ρ* rank-order correlations among attitude variables. For our third objective, we used Spearman’s rho to measure rank-order correlation between independent variables and the dependent variable (future compliance with the pacu size rule). We report the ordered Probit regression model to describe the effect of trust in scientists, specifically, on frequency of noncompliance, and we speculate on why certain demographic variables also correlate with noncompliance rate. Data were analyzed in R3.4.4 (57).

## Results and discussion

Forty-one respondents agreed to respond to be interviewed and three people refused to participate, resulting in a response rate of 93.2 percent. Of the respondents, the majority were men with two or more children, and fewer than half had finished primary school. Education level was inversely correlated with age (*r =* −0.46) and years fishing (*r =* −0.28). On average, participants were over 48 years old with more than 38 years of fishing experience. Most participants were unable to estimate their monthly income but all use fishing as their primary work during the open fishing season (March through September or October). During the closed season, they earned a monthly stipend from the government that was slightly above minimum wage. Although all participants lived within a 50-km radius of a scientific research station and were aware of scientist’ work, few had interacted previously with scientists conducting research on the fisheries in the region (Table 1).

**Table 1.** Demographics of participating *in-loco* professional fishermen in one fishing community in the Pantanal.

| Participant Information |  |
| --- | --- |
| Gender | 34 Men, 7 Women |
| Level of Education<sup>a</sup> | Mean 0.85, sd 1.06 |
| Age | Mean 48.09, sd 13.70 |
| Years Fishing | Mean 38.44, sd 17.45 |
| Number of Children | Mean 3.05, sd 2.44 |
| Previous interaction with environmental police? | 38 Yes, 3 No |
| Previous interaction with a scientist? | 5 Yes, 36 No |

<sup>a</sup> Levels of education: 0 = no education; 1 = some primary school; 2 = completed primary school; 3 = completed secondary school; 4 = completed tertiary education.

### 1. Fisheries noncompliance rates

Our first objective focused on assessing rates of noncompliance. We asked all study participants (n = 41) two questions about rates of compliance to assess views on the frequency of noncompliance in the community. Participants reported violations occurring frequently in the community. A majority (n = 25, 60.9%) agreed or agreed strongly that violations were common. In commentaries, participants accused amateur fishers without fishing licenses of violating the law the most. Most reported that they, personally, were usually compliant with the pacu catch size rule (34 or 85% indicated they would break the rule “sometimes,” “rarely,” or “never” in the coming year). A small minority of participants indicated they would break the rule all the time (n = 4, 9.7%); three participants reported breaking the rules often (7.3%) and six said they never would break the rules (14.6%). The average rate of self-reported future noncompliance among participants was 2.5, or between “rarely” and “sometimes.” The sentiment in the community of professional fishermen is that there are others—amateurs, professionals, and tourists—violating the fishing rules often, but virtually nobody identified themselves as part of the problem. Noncompliance rates correlated negatively with age (*ρ* = 0.22) and positively with education level (*ρ* = 0.37), which in turn correlated negatively with each other (*ρ* = 0.50). Older, less educated people tended to comply with laws with a greater frequency than younger, more educated people; this is in accordance with other studies that have found age to be a significant factor in determining compliance (58).

Our survey questions related to noncompliance were projective and prospective (asking about others’ noncompliance and estimates of future noncompliance) to protect respondents from potential legal consequences of reporting their own past or present rule breaking. However, the idea that noncompliance with regulations is prevalent in the Pantanal is not particularly controversial, nor is the behavior particularly covert. This observation was amply supported by anecdotal evidence from community members and personal experiences by the lead author during the data collection period. Accounts contradicting the notion that noncompliance is prevalent and overt tend to focus on more severe forms of rule breaking (e.g., using nets to catch hundreds of pounds more than the permitted weight) compared with the relatively small violation on which we focused here (50). For example, the majority of undersized fish we observed were still adult fish, just not quite large enough to meet the size minima prescribed by law. This contrasts with violators who were intentionally fishing dourado, a protected species fish for which fishing is banned, for weeks at a time (observed in 2017), and with others who use fishing nets (observed in 2016) and dynamite (anecdote in 2016 in a different region of the Pantanal). Respondents, in their comments, highlighted these differences between their own noncompliance and the noncompliance of those who were truly damaging the environment, and frequently attributed the behavior of others to inherent bad character. Their comments provide evidence for the fundamental attribution error (59,60), which could suggest that due to correspondence bias people attribute their own behavior to external factors whereas behavior of others reflects internal flaws. This error has been shown to be a factor in environmental decisions of hunters and may be relevant in fishermen as well (61). Additional research would benefit this discourse.

Regardless of motive, the noncompliance rates studied here may or may not cause extreme environmental harm. The idea that more severe forms of noncompliance may be viewed differently is one that is also in keeping with the idea that professional fishermen have only a nominal negative impact on the environment. According to (62), overfishing is one of a bevy of factors causing harm in the Pantanal, and possibly less important when compared with environmental damage produced by sewage and other pollution, climatic changes, and damming of upstream tributaries. Even if instances of noncompliance are commonplace in the community of professional fishermen, it does not necessarily mean that they are the instigators of widespread environmental damage to the Pantanal. However, local people’s cooperation with managers is necessary for successful management of the resource.

### 2. Community perceptions of risk and management

A range of motivations were presented as underlying noncompliance with fisheries rules, including lack of enforcement (n = 29, 70.7% agreed or agreed strongly that it was a factor) and lack of knowledge of rules (n = 3, 7.3% agreed or agreed strongly). When individuals were asked about their attitudes, most generally seemed aware of environmental problems and risks (Table 2). Many negatively viewed aspects of the management structure and the procedural fairness in the region; however, most disagreed that the management agency was actively deceiving them.

**Table 2.**
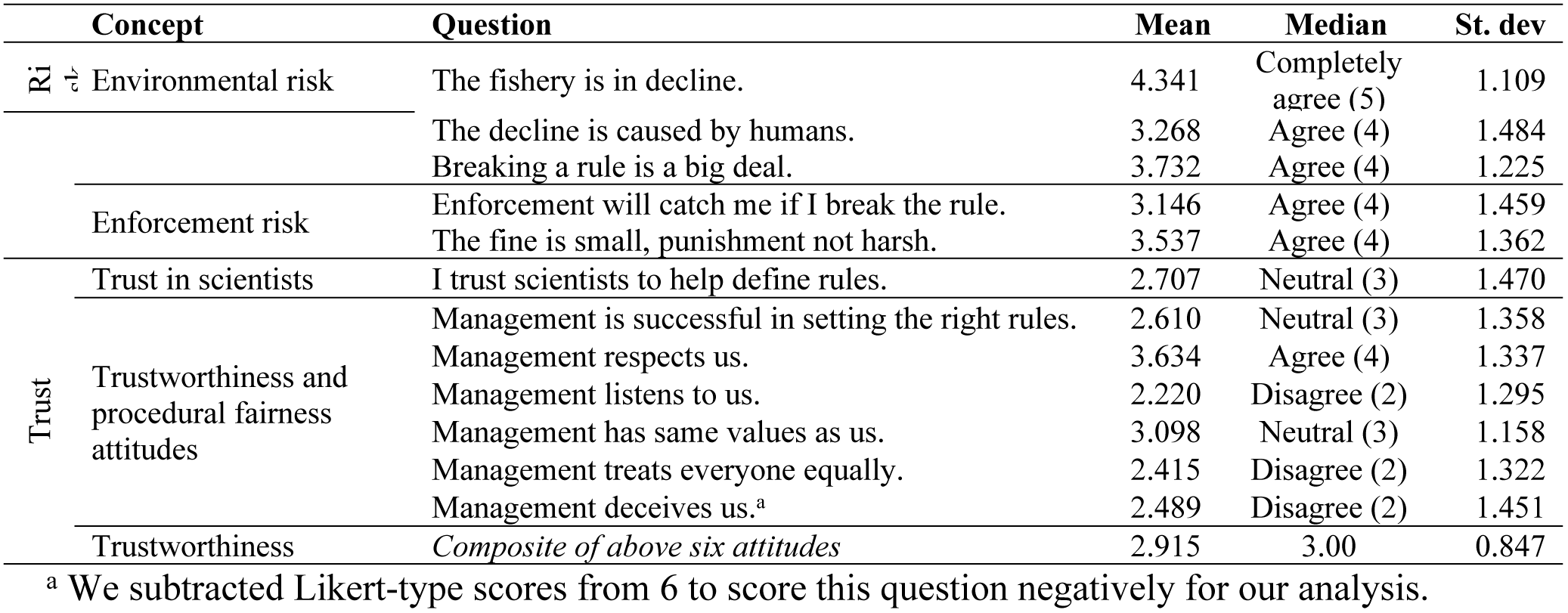
Means, medians, and standard deviation of responses to Likert-type attitude questions focused on noncompliance with fisheries rules in the Brazilian Pantanal, April-August 2016.

General attitudes about environmental risks among respondents in this study indicated an interest in the environment and its conservation. The majority of environmental attitude questions we asked respondents to provide are derived from those in the literature, and responses from a community that depends on natural resources for its livelihoods and subsistence is not surprising. Some individuals, however, indicated skepticism about whether humans are the ones causing environmental harm. This attitude was not correlated with any others but is notable— many who said the fishery is in a decline then suggested that it was primarily caused by the increasing population of piscivorous species such as the giant river otter (*Pteroneura brasiliensis*) and caiman (*Caiman yacare*). These species have been recovering from decimation in the late-20th century due to the pelt trade and are much more abundant than they were merely decades ago. Scientists tend to reject the contention that the recovery of predator populations has adversely affected the fishery, instead suggesting that healthier predator populations may actually protect fish stocks (63).

Respondents’ attitudes about the natural resource management agency portrayed the institution in a mixed light. Although very few claimed that the managers were actively deceiving them, almost none seemed to think that they had sufficient voice to influence rules. Trust Trustworthiness and procedural fairness attitudes This is noteworthy given management in the Pantanal is designated as a co-management system—one in which stakeholders contribute to rulemaking decisions. However, the ability to contribute to the rulemaking decisions is limited to those who can travel some 50km to participate in fishermen’s colony meetings, or some 150km to participate in CEPESCA meetings. Furthermore, respondents augmented their responses about enforcement with anecdotes of how police sometimes invade their homes without a warrant. Views of risk of being caught for a violation were mixed—although a majority agreed or agreed completely (n = 24, 58.5%) that if they violated a rule they would be caught, a majority (n = 26, 63.4%) also said that the penalty was relatively small. It was therefore unclear what sort of deterrent effect enforcement had in this region; however, age correlated positively with perceived chance of getting caught (*ρ* = 0.52) and negatively with the penalty being small (*ρ* = −0.24). This follows the logic of other noncompliance studies suggesting that older respondents are more risk averse (16).

In a situation like that in the Pantanal, wherein one commission consists of enforcers, researchers, and legislators, individuals’ views on the structure as a whole may depend on interactions with different parts. We measured trust by analyzing agency trustworthiness as well as asking about trust in scientists directly in an effort to differentiate between scientists and the rest of the management agency. This trust inquiry is not without complications. Trust has been defined in the literature as a function of trustworthiness and risk (37). Questions about procedural fairness and whether people have a voice in making rules are among those that are considered part of institutional trustworthiness in the literature. Trust, in Portuguese, is the same word as confidence (*confiança*), although some authors in the English literature have stressed the differences between these two constructs (64). In our study, trust in scientists correlated moderately (*ρ* = 0.49) with our agency trustworthiness composite score and with perceived success in the agency setting the rules (*ρ* = 0.34). The success question comes from the *ability* subset of trustworthiness questions (37), as one would expect—people who trust the science behind rules trust the ability of the organization to set the right rules. All of these trust variables correlated positively with age and negatively with education level, bringing into question the reason for apparent less trust in scientists by the more educated in the community. The questions asked about institutional trust did not differentiate well between the different roles the agency plays—in addition to scientists, there are politicians and enforcement officers in CEPESCA, all of whom play a part in rulemaking, but none of whom singularly control the creation of each rule. It is possible that distrust in one group could be projected onto another group within the management structure. This is possibly the reason trust in scientists correlates only moderately with trust in the management agency as a whole. The questions did not consider the interactions people may have had with enforcement officers and how those interactions might have shaped trust in other agency members. Trust in scientists was also not differentiated here from trust in science, itself (19), which some studies have found to be in decline (65). Future research could differentiate between trust in science, trust in scientists, trust in police, trust in rule makers, and trustworthiness of the agency as both a rule maker and a rule enforcer.

### 3. Factors contributing to individual noncompliance rates

We focused our questions about the future noncompliance of the pacu size rule specifically to measure people’s reasons for noncompliance. Level of education correlated positively with rate of noncompliance with this rule, and age and years fishing correlated negatively. Level of education also correlated negatively with trust and trustworthiness, while age and years fishing correlated positively with trust and trustworthiness. Other authors have speculated that older people tended to comply more because they are more risk-averse and more involved in management decisions. Among the attitudes related to environmental risk, enforcement risk, management trustworthiness, and procedural fairness, only two measures significantly predicted frequency of noncompliance—trust in scientists to help define rules and the composite trustworthiness. Trust in scientists is the most predictive in an ordered Probit regression model (pseudo-R^2^ 0.241) of frequency of noncompliance. Multivariate models including enforcement risk as an alternative and independent factor in noncompliance did not return significant results; other univariate models with age and education level were less significant and far less predictive (pseudo-R^2^ < 0.05) than trust and trustworthiness models. Nonparametric rank correlation tests returned similar results, with P < 0.001 and a particularly high negative correlation between trust in scientists and noncompliance rates (*ρ* = −0.717).

That respondents’ trust in scientists affected stated rates of noncompliance with a rule influenced by empirical evidence reflects with parallel conclusions in the literature. Trust in management more generally, both in the form of procedural justice (38) and in institutional trust (34,42) has been shown in the literature to be related to compliance, although these studies focus on management as a whole, as opposed to researchers specifically. Trust in science and scientists also logically may be related to understanding of research, something that could in turn be related to education and age, depending on how educational opportunity has evolved through time. Age in this study was correlated with education level and years fishing, which were more reliable predictors of noncompliance rates. We found no evidence that enforcement risk or risk aversion play a part in compliance decisions in this region. Although in other contexts authors have argued that age is related to compliance because older people are more risk averse, in this context it appears that age may be related to risk aversion and trust in management, but that only trust in management is predictive of noncompliance.

## Implications for natural resource management and conclusions

This study set out to explore the human cognitions and behaviors underlying inland IUU fishing. Because rules exist to help mitigate risks associated with human pressure on the environment, decreasing rates of rule noncompliance can help maximize rule effect. We explored noncompliance in a context where compliance has understudied. We focused on attitudes that are rarely examined in a freshwater context, but which could be especially important to voluntary compliance due to remoteness and difficulty of enforcement. Although our study context was unique, it embodies conditions common in other key biodiversity areas around the world. Below, we discuss the implications of our findings for conservation criminology theory as well as the effective practice of natural resource management.

We found that in one community of Pantanal professional fishermen, noncompliance was overt and commonplace, a fact that we personally observed on many occasions. Although aspects like enforcement, procedural justice, and environmental risk can be important, the most important factor influencing noncompliance rate among the population of professional fishermen in this study group in the Pantanal was trust in the scientists helping to define the rules. Each violation of a rule is an example of IUU fishing, and although each violation may individually be small, the collective effect of violations can be large. There may be collateral effects of “small” transgressions of the rules, such as the promotion of a culture of violating rules and the lack of cooperation with enforcement and legislators to catch larger violators and write better rules. The exact amount of damage that violators of fishing rules cause is an empirical question not addressed in this study. Thus, reducing all types and sizes of IUU fishing bears merit. Trust in scientists was a predictive factor for noncompliance decisions in our study community of fishermen in the Brazilian Pantanal. Increasing trust in scientists may be one mechanism for decreasing rates of noncompliance among our study population.

Building trust is known to be challenging. Davenport et al. (42) showed that in spite of clear indications that trust in management is necessary for success, a number of barriers exist to building trust, including lack of community engagement, knowledge gaps, and competing values. Many of these barriers appeared present in our study community. Very few of this study’s participants had interacted with scientists in the past, potentially explaining a lack of mutual understanding and mismatching values. Rudolph & Riley (35) argued that gains in trust may be possible through changes in structure of procedural justice of the management system. Encouraging community members to share their voice can be critical for the success in a co-management system, and the fact that so many people in our case study group feel that management did not listen to their views highlights one opportunity for potential improvement. It is possible that more effective community engagement by scientists could help advance community members’ understanding about their participatory rights in the management structure. This in turn might amplify positive perceptions of procedural justice of managers in the community. Future research would help explore these ideas further.

Trust in scientists is unlikely to be the primary driver of noncompliance decisions in every natural resource management system—our results are specific to one community in the Pantanal. However, a confluence of considerations from the case study group and Brazilian Pantanal may help explain the conditions under which trust in scientists may be more important than other factors. First, the community of professional fishermen in the Pantanal is not unlike communities around the world in key biodiversity areas; it historically has had little access to education and there is a rift between the scientific elites doing research and creating laws and the local population. The extant literature demonstrates the value of using local people’s knowledge and understanding of biological systems to improve the quality of scientific research in general (66), detailing a slew of specific benefits (67) for conservation worldwide (68) and in Brazilian fisheries in particular (69). The prolific influence of trust in scientists on frequency of noncompliance in this Pantanal community further underlines a different advantage of closing the gap of understanding between scientists and locals—that it may also result in more favorable conservation outcomes because of more consistent and widespread compliance with environmental rules.

## Acknowledgments

We thank the Fulbright Scholars Program and all of those who contributed to the theoretical formulation of this project, including colleagues at Michigan State University and the Federal University of Mato Grosso, as well as the Pantanal fishermen who made this work enjoyable and whose rights and nature this work seeks to protect.

